# Low-intensity focused ultrasound pulsation along the anterior–posterior thalamic axis differentially modulates the latency of reporting conscious visual experience

**DOI:** 10.64898/2026.08.21.746115

**Authors:** Hyunwoo Jang, Jiyang Liu, Anthony G. Hudetz, Zirui Huang

## Abstract

**Background:** Transcranial low-intensity focused ultrasound (LIFU) neuromodulation can alter human task performance depending on target and acoustic configurations. However, single behavioral endpoints cannot locate effects within multistep tasks, and predefined target labels ignore acoustic variations.

**Objective:** To determine whether thalamic LIFU affects visual categorization or subsequent subjective report latency and whether the effects vary with target, acoustic parameter, and beam location.

**Methods:** Sixty healthy adults were randomized to 70% or 5% duty cycle (DC70 or DC5) and received sonication on four left thalamic targets with matched pulse repetition frequency (10 Hz) and temporal-average intensity (0.72 W/cm^2^). Behavioral models tested target-by-DC interactions in categorization (RT1) and subjective report (RT2) latencies. Spatial analyses correlated focal spot coordinates and voxel-wise intensity from 179 acoustic simulations to baseline-adjusted RT2.

**Results:** The target-by-DC interaction was detected for RT2 but not RT1. At the ventroposterior thalamic target, adjusted RT2 was 55.9 ms longer under DC70 than DC5. More anterior focal spots shortened RT2 under DC70 but increased RT2 under DC5. Correlation between intensity and adjusted RT2 significantly differed between DC70 and DC5 in 18.8% of thalamic voxels. These voxels formed an anterior mediodorsal–motor set and a posterior pulvinar-dominant set.

**Conclusions:** The latency of reporting conscious visual experience, but not categorization latency, was affected by thalamic LIFU. This effect varied jointly with anterior–posterior target engagement and acoustic configuration. Analyzing sequential reaction times separately and treating field variation as an anatomical variable revealed associations not fully captured by a single endpoint or predefined target labels.

**HIGHLIGHTS:**

- Ultrasound changed conscious visual report latency, not categorization latency.
- Reports were slower at 70% than 5% duty cycle in ventroposterior thalamus.
- Anterior sonication made reports faster at 70% and slower at 5% duty cycle.
- Posterior sonication made reports slower at 70% and faster at 5% duty cycle.

## Introduction

Low-intensity focused ultrasound (LIFU) enables noninvasive perturbation of cortical and deep brain structures with relatively high spatial precision [1]. Human studies have examined the neuromodulatory effects of LIFU on perception, inhibitory control, reward processing, emotional evaluation, attention, and memory [2–7]. However, behavioral effects are often summarized by a single endpoint, such as accuracy, perceptual sensitivity, or total reaction time. In a task requiring sequential responses, a single endpoint cannot show whether a stimulation-evoked difference arises during the initial decision or a later report. Measuring the latency of each response separately can locate the stimulation effect within the task and constrain its functional interpretation.

The thalamus is an important target for neuromodulation because its anatomically and functionally distinct subdivisions connect with cortical networks involved in sensory processing, perceptual evaluation, and motor execution [8–12]. Human recordings and electrical stimulation have shown that thalamic activity can predict or alter reaction time [13,14]. LIFU studies have also reported reaction time changes in visual motion and attention tasks [15,16]. However, whether thalamic sonication affects successive response intervals differently within a multi-step perceptual task remains unclear.

We previously investigated thalamic LIFU during a near-threshold visual task in which participants first categorized an image and then reported whether they had seen it [17]. The study tested four thalamic targets and two active acoustic configurations that used 70% or 5% duty cycle (DC70 or DC5), with matched pulse repetition frequency (PRF) and temporal-average intensity. Visual recognition sensitivity differed across targets and two acoustic configurations [17], but this study did not investigate whether LIFU effects were expressed in the categorization latency, the subsequent subjective report latency, or both.

The previous analysis also treated variation across simulated acoustic fields as an anatomical predictor rather than relying only on target labels. We revealed that sensitivity change was associated with the cell composition within the focal spot of each simulated field [17]. Whether this approach can be extended to reaction time analysis remains unknown.

Here, we reanalyze our published dataset to determine whether thalamic LIFU neuromodulation altered the reaction times of object categorization (RT1) and subsequent report of visual experience (RT2). We first test whether thalamic target and acoustic protocol (i.e., DC70 and DC5) interact for each RT. We then determine whether reaction time varies with the focal spot position along the left–right, anterior–posterior, or dorsal–ventral axis. Finally, we ask whether simulated acoustic intensity at each thalamic voxel is associated with the reaction time, and whether this association differs between DC70 and DC5. We hypothesize that reaction time differences between the two protocols depend on thalamic target and that simulated acoustic fields reveal spatial variation not captured by assigned target labels.

## RESULTS

### EXPERIMENTAL DESIGN

This study used two active acoustic configurations, DC70 and DC5, with matched temporal-average intensity and pulse repetition frequency (PRF) (Fig. 1). Sixty subjects were randomized to two arms, DC70 or DC5, with 30 subjects assigned to each configuration. Six subjects were excluded due to technical issues (see Methods: Subjects) and 54 subjects remained for analysis (27 per arm). Each subject completed an initial baseline block, four LIFU-ON blocks, and a final baseline block. During the LIFU-ON blocks, four regions of the left thalamus were targeted in pseudo-randomized and counterbalanced order: ventroanterior, ventroposterior, dorsoanterior, and dorsoposterior thalamus (VA, VP, DA, and DP) in Tian Subcortical Atlas Scale-II [8].

**Fig. 1:**
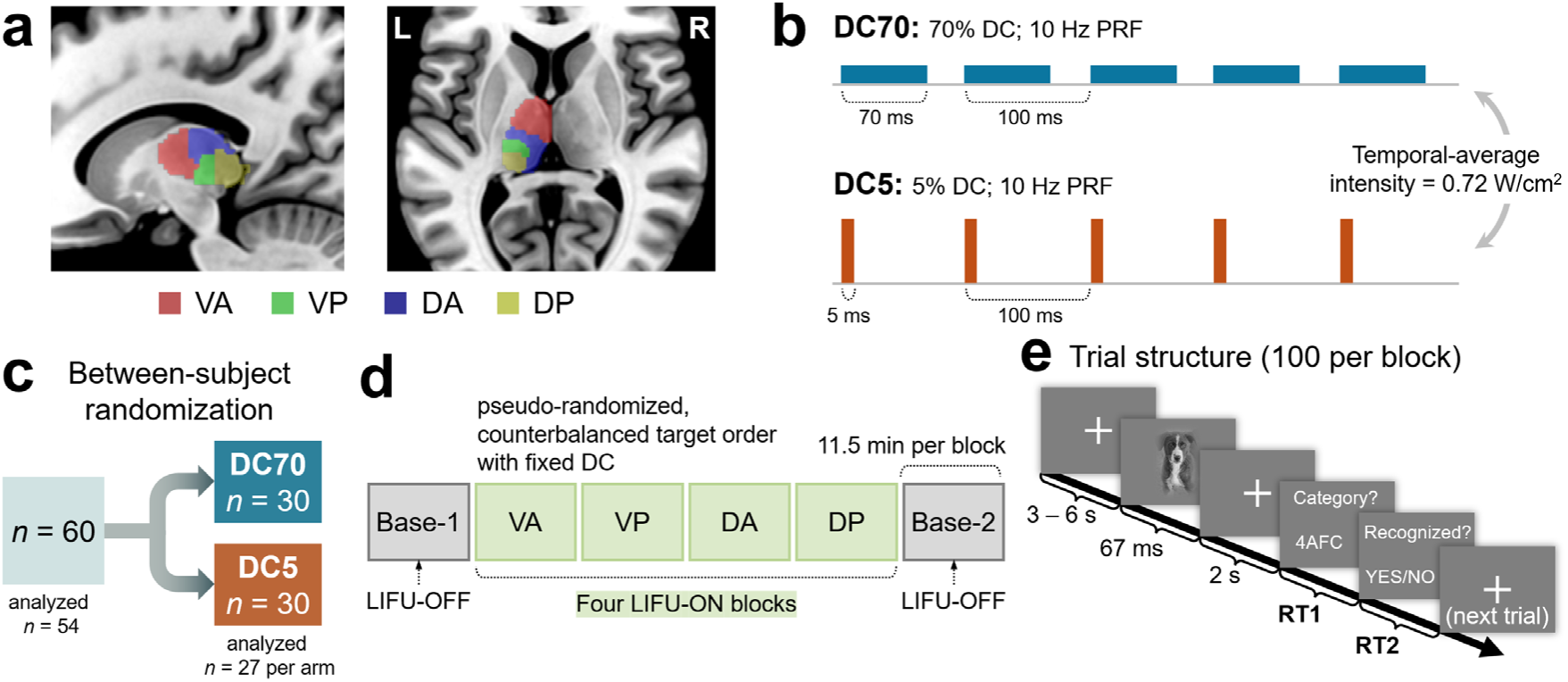
Experimental design and acoustic configurations. (a) Four target regions in the left thalamus. (b) Two acoustic configurations used in the study. DC70 used a 70-ms pulse and DC5 used a 5-ms pulse within each 100-ms cycle. Pulse repetition frequency (PRF) and derated spatial-peak temporal-average intensity were matched at 10 Hz and 0.72 W/cm^2^, respectively. (c) Sixty subjects were randomized between DC conditions, and 54 were analyzed. (d) Each subject completed six blocks, including Baseline-1, four LIFU-ON blocks, and Baseline-2. During LIFU-ON blocks, four targets were sonicated in pseudo-randomized and counterbalanced order. (e) Trial structure. RT1 and RT2 were the latency from the onset of the prompts to the respective response. 4AFC, four-alternative forced choice; DA, dorsoanterior thalamus; DC, duty cycle; DP, dorsoposterior thalamus; RT, reaction time; VA, ventroanterior thalamus; VP, ventroposterior thalamus.

Each block contained 100 trials. On each trial, subjects were first asked to categorize the near-threshold image in a four-alternative forced choice task (Q1) and then to report whether they had a meaningful visual experience of it (Q2). RT1 was defined as the latency from the onset of the Q1 prompt to the response. Q2 prompt was displayed immediately after Q1 response, and RT2 was defined as the latency from the onset of the Q2 prompt to the response. Each reaction time outcome was then summarized as the median within each block. Because reaction time declined over the first trials as the subjects adapted to the task, trials 1–10 were excluded from median calculation (Fig. S1).

### Thalamic target and duty cycle interact on reaction time to report conscious visual experience

We fit block-level linear mixed effects (LME) models for RT1 and RT2 during the 209 LIFU-ON blocks (DC70: *n* = 103; DC5: *n* = 106). Each model included target, DC, their interaction, the corresponding Base-1 outcome, sonication order, and four other nuisance variables. The models also included subject-specific random intercepts and slopes. Satterthwaite *F* tests detected a target-by-DC interaction for RT2 after false discovery rate (FDR) correction (*F* = 4.258, *p*_FDR_ = 0.0132), but not for RT1 (*F* = 0.730, *p*_FDR_ = 0.5361) (Fig. 2, Table S1).

**Fig. 2:**
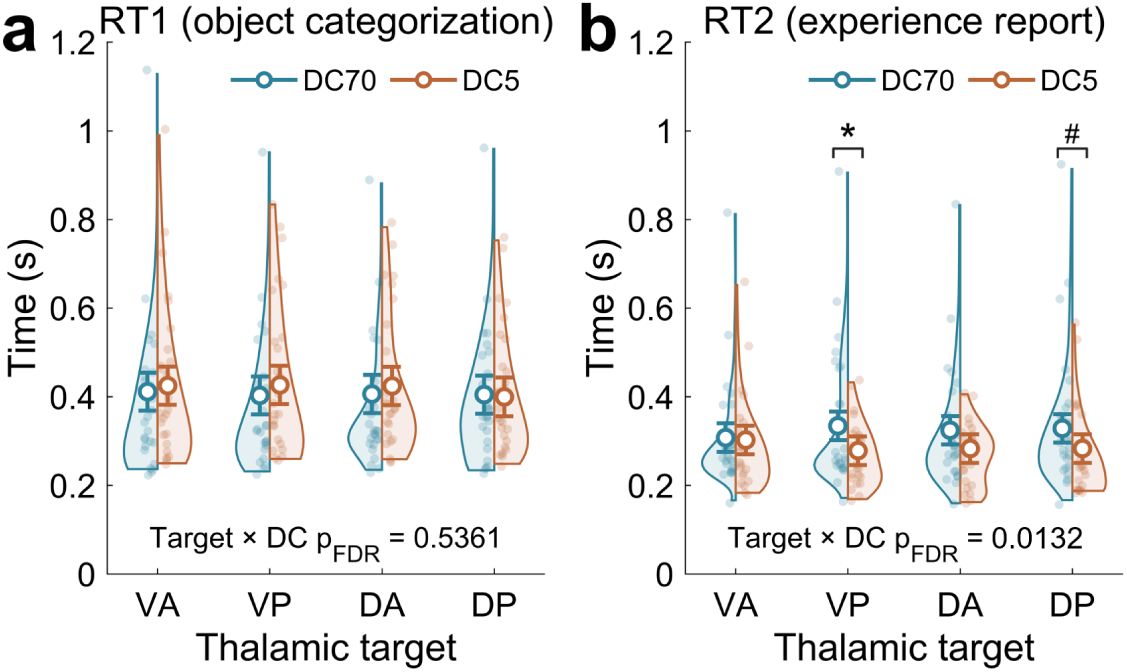
Region-level effect of LIFU on reaction time. Reaction time distributions for (a) RT1 for object categorization and (b) RT2 for conscious visual experience report. Violins show unadjusted distributions of block medians for DC70 and DC5. Omnibus target-by-DC *p*-values were FDR-corrected across RT1 and RT2. Within RT2, target-specific DC70-minus-DC5 contrasts were FDR-corrected across the four targets. An asterisk denotes *p*_FDR_ < 0.05 and a pound indicates uncorrected *p* < 0.05. Open circles and error bars show model-estimated marginal means and 95% confidence intervals. DA, dorsoanterior thalamus; DC, duty cycle; DP, dorsoposterior thalamus; RT, reaction time; VA, ventroanterior thalamus; VP, ventroposterior thalamus.

Post-hoc contrasts were restricted to RT2 and corrected across the four targets. RT2 was 55.9 ms longer under DC70 than DC5 at VP (95% CI, 14.5–97.3; *p*_FDR_ = 0.0353). DP showed a nominal difference of 45.5 ms (4.3–86.7; *p*_FDR_ = 0.0616, uncorrected *p* = 0.0308). The contrasts at DA and VA were not significant (Fig. 2b, Table S2).

### Anterior–posterior position of the focal spot affects reaction time oppositely across duty cycles

We simulated the acoustic fields from 179 LIFU-ON blocks of 47 subjects. For each field, the focal spot was defined as the MNI coordinate of maximum simulated intensity. The standard deviation of focal spot location was 6.7 mm along the anterior–posterior axis, 6.3 mm along the left–right axis, and 5.2 mm along the dorsal–ventral axis.

We then correlated MNI coordinates of the simulated focal spots with RT2 residuals (i.e., baseline-adjusted RT2) from a separate LME model excluding target and DC. Under DC5, more posterior beams were associated with shorter baseline-adjusted RT2 (*ρ*_DC5_ = 0.2847). This association reversed under DC70 (*ρ*_DC70_ = –0.1830), and this between-DC contrast was significant (Δ Fisher-*z* = –0.4779, *p*_FDR_ = 0.0141; Fig. 3b). No corresponding contrast survived correction along the left–right or dorsal–ventral axis (*p*_FDR_ > 0.05; Fig. 3a,c, Table S3). RT1 did not show contrast in any of the three axes (Fig. S2).

**Fig. 3:**
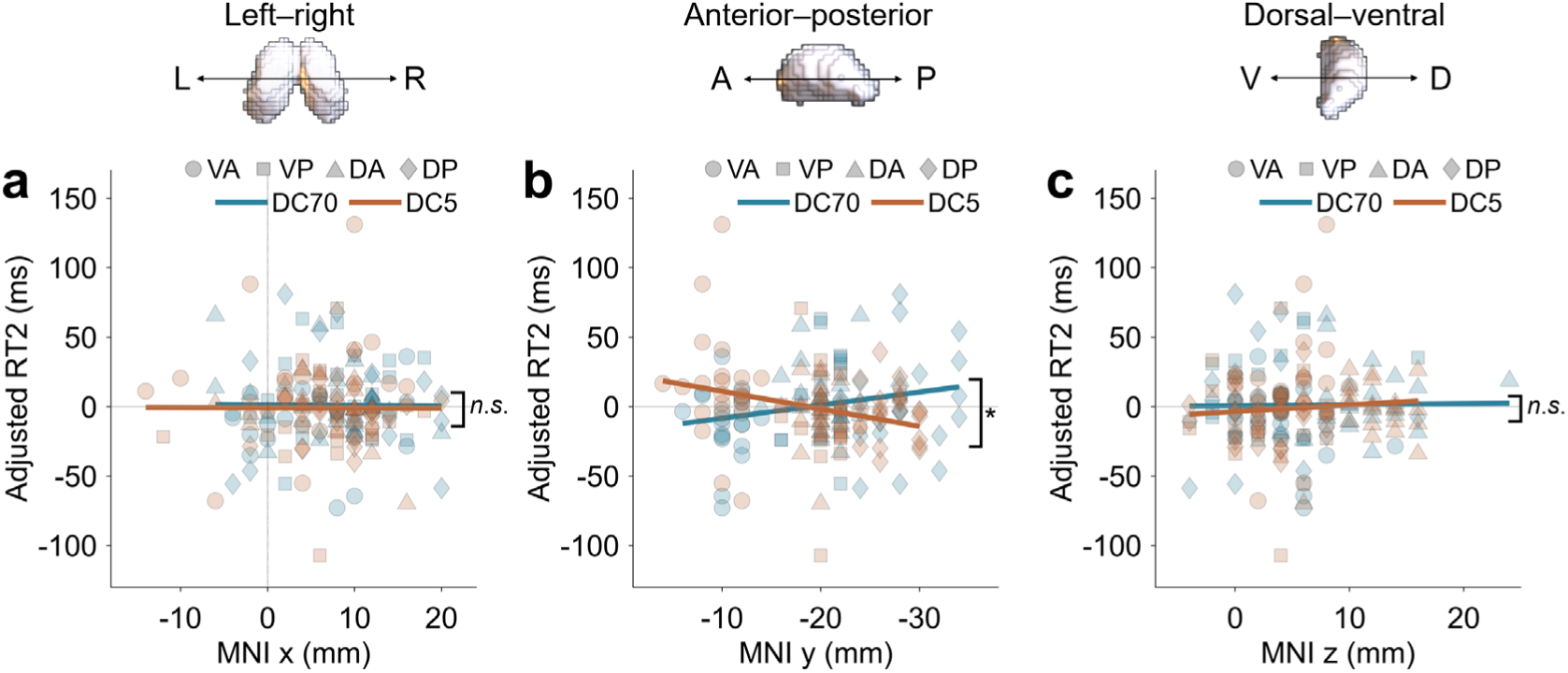
Correlation between focal spot location and baseline-adjusted RT2. The MNI coordinates of field-wise focal spot along the (a) left–right, (b) anterior–posterior, and (c) dorsal–ventral axes are plotted against baseline-adjusted RT2. Focal spot was defined as the voxel showing highest simulated intensity. Teal and orange lines are linear fits provided only for visualization. Symbol shapes indicate intended target regions. An asterisk marks the between-DC difference that survived FDR correction across the three axes. In (a), vertical dash denotes midsagittal line (*x* = 0 mm). Sample sizes are *n* = 84 from 22 subjects for DC70 and *n* = 95 from 25 subjects for DC5. DA, dorsoanterior thalamus; DC, duty cycle; DP, dorsoposterior thalamus; RT, reaction time; VA, ventroanterior thalamus; VP, ventroposterior thalamus.

Adjusted RT2 is relative to its covariate-controlled prediction and should not be directly interpreted as acceleration or deceleration of RT2 in absolute terms. What this result demonstrates is that predicted RT2 associated with the anterior–posterior position of the acoustic beam, and its trend significantly differed between acoustic configurations.

### Associations between acoustic intensity and reaction time reverse along the anterior– posterior thalamic axis

We mapped the spatial trend of LIFU effects on RT2 across bilateral thalamic voxels (Fig. 4a). Analysis mask was defined as the union of bilateral thalamus in Tian Subcortical and Morel atlases [8,18,19], and 1,923 native 2-mm isotropic voxels fully contained within the mask was investigated. At each voxel, Spearman correlation between simulated acoustic intensity and baseline-adjusted RT2 was calculated separately under DC70 and DC5 and was transformed to Fisher-*z*, denoted *z*_DC70_ and *z*_DC5_, respectively. Therefore, positive *z*_DC70_ or *z*_DC5_ at a given voxel implies longer adjusted RT2 when it receives higher acoustic intensity.

Anterior and posterior regions showed opposing trends (Fig. 4b). *z*_DC70_ values were negative at anterior regions and positive at posterior regions. This pattern was reversed at DC5, with positive *z*_DC5_ values anteriorly and negative values posteriorly. The anterior–posterior slope was significant for DC5 (*p*_FDR_ = 0.0216; Table S4) but not DC70 (*p*_FDR_ > 0.05). Thus, DC5 sonication was associated with RT2 differently at anterior and posterior thalamus. RT1 did not show a significant anterior–posterior slope (Fig. S3a).

**Fig. 4:**
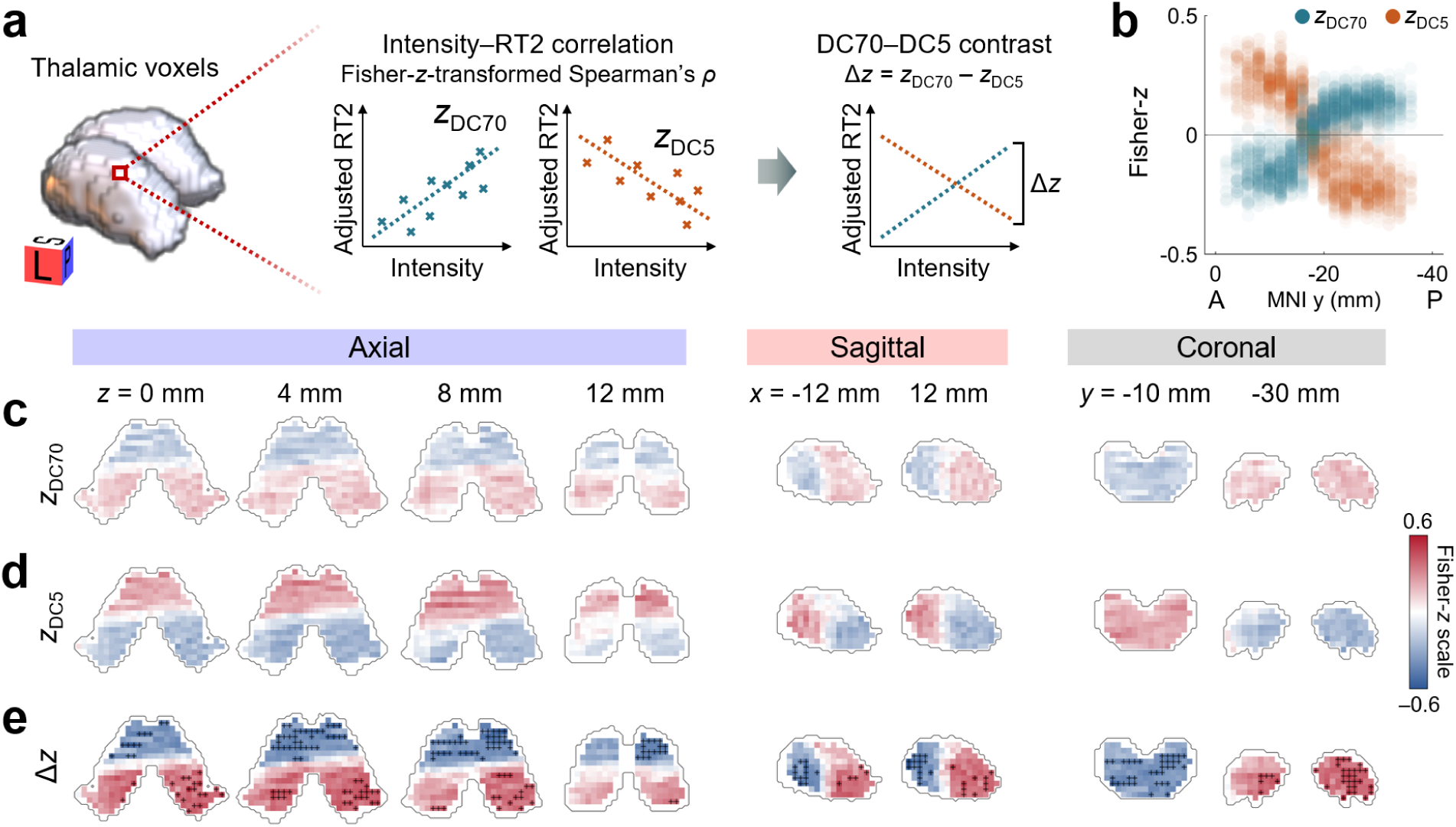
Voxel-wise associations between acoustic intensity and baseline-adjusted RT2. (a) Schematic of voxel-wise analysis. At each voxel, simulated acoustic intensity was Spearman-correlated with baseline-adjusted RT2 separately under DC70 and DC5. Correlations were Fisher-*z* transformed to yield *z*_DC70_ and *z*_DC5_, and the between-DC contrast was defined as Δ*z* = *z*_DC70_ – *z*_DC5_. (b) Values of z_DC70_ and *z*_DC5_ plotted against the MNI *y* coordinate of 1,923 analyzed voxels. (c-e) Unthresholded maps of *z*_DC70_, *z*_DC5_, and Δ*z* shown in axial sections at *z* = 0, 4, 8, and 12 mm, sagittal sections at *x* = −12 and 12 mm, and coronal sections at *y* = −10 and −30 mm. Outlines indicate boundary of the Tian S2–Morel union thalamic mask. Black crosses in (e) highlight voxels that survived FDR correction. DC, duty cycle; RT, reaction time.

We then tested the between-DC contrast by calculating Δ*z* = *z*_DC70_ – *z*_DC5_ at each voxel. This can reveal voxels at which the effect of LIFU significantly differs across the two DC conditions. There was a significant Δ*z* slope along the anterior–posterior axis (*p*_FDR_ = 0.0216; Table S4). Subject-level permutations tested the significance of Δ*z* contrast at each voxel. After FDR correction, 362 voxels (18.8%) survived. These comprised 216 voxels with negative Δ*z* and 146 voxels with positive Δ*z* (Fig. 4e). The negative-Δ*z* voxel set occupied anterior positions (MNI *y*, −16 to −2 mm), whereas the positive-Δ*z* voxel set resided posteriorly (−36 to −20 mm), with no overlap in their distributions (Fig. S4b). All negative-Δ*z* voxels showed negative correlations under DC70 and positive correlations under DC5. Conversely, all positive-Δ*z* voxels showed positive correlations under DC70 and negative correlations under DC5. Thus, every FDR-surviving voxel showed a reversal in the direction of the intensity–RT2 association between DC protocols. The direction of these associations is summarized in Table 1.

**Table 1:** DC-dependent changes in RT2 at the anterior and posterior voxel sets.

|  | Greater intensity at DC70 | Greater intensity at DC5 |
| --- | --- | --- |
| Anterior, negative-$\Delta z$ voxel set | Shorter RT2 | Longer RT2 |
| Posterior, positive-$\Delta z$ voxel set | Longer RT2 | Shorter RT2 |

Entries indicate the direction of the association between simulated acoustic intensity and baseline-adjusted RT2.

To characterize their anatomical distribution, we measured how much of each voxel set fell within Tian Scale-III parcels and Morel nuclei (Fig. 5). In the Tian atlas, the anterior negative-Δ*z* voxel set was almost entirely VA (99.0%). The posterior positive-Δ*z* voxel set was concentrated in DP (56.2%), with additional contributions from VP (21.4%) and DA (11.9%). In the Morel atlas, the anterior voxel set was distributed across lateral and medial nuclei, with the largest contributions from the parvocellular mediodorsal nucleus (19.8%), ventral division of ventrolateral posterior nucleus (16.5%), and parvocellular ventral anterior nucleus (14.1%). The posterior voxel set was concentrated in posterior nuclei, particularly the medial pulvinar (60.4%) and lateral pulvinar (8.6%).

**Fig. 5:**
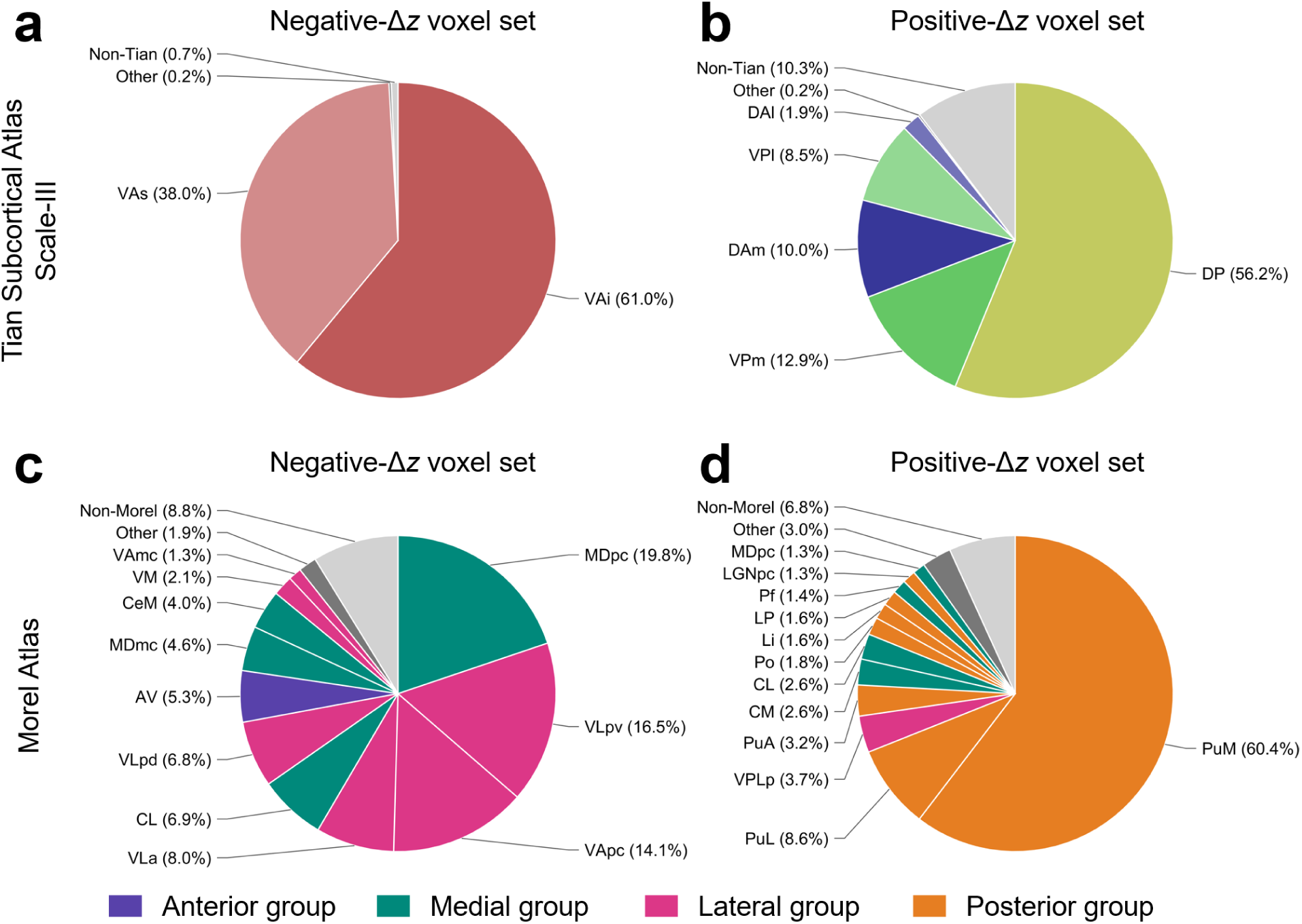
Anatomical composition of voxel sets showing significant between-DC difference. Pie charts showing the distributions of the anterior negative-Δ*z* (216 voxels) and posterior positive-Δ*z* (146 voxels) voxel sets across (a,b) the Tian Scale-III atlas and (c,d) Morel atlas. Percentages indicate each region’s share of the voxel set’s total volume. Atlas regions contributing less than 1% were combined as Other; volumes outside each atlas are labeled non-Tian or non-Morel. Morel colors indicate anterior, medial, lateral, and posterior nuclear groups. Tian abbreviations: DAl, lateral dorsoanterior thalamus; DAm, medial dorsoanterior thalamus; DP, dorsoposterior thalamus; VAi, inferior ventroanterior thalamus; VAs, superior ventroanterior thalamus; VPl, lateral ventroposterior thalamus; VPm, medial ventroposterior thalamus. Morel abbreviations: AV, anteroventral nucleus; CeM, central medial nucleus; CL, central lateral nucleus; CM, centromedian nucleus; LGNpc, parvocellular lateral geniculate nucleus; Li, limitans nucleus; LP, lateral posterior nucleus; MDmc, magnocellular mediodorsal nucleus; MDpc, parvocellular mediodorsal nucleus; Pf, parafascicular nucleus; Po, posterior nucleus; PuA, anterior pulvinar; PuL, lateral pulvinar; PuM, medial pulvinar; VAmc, magnocellular ventral anterior nucleus; VApc, parvocellular ventral anterior nucleus; VLa, ventral lateral anterior nucleus; VLpd, ventral lateral posterior nucleus, dorsal division; VLpv, ventral lateral posterior nucleus, ventral division; VM, ventromedial nucleus; VPLp, ventral posterior lateral nucleus, posterior division.

## DISCUSSION

Previous LIFU studies indicate that reaction time effects can be selective to trial or response type [15,20]. Assessing separate reaction times for sequential phases of a perceptual task provides a more complete characterization of the effect of neuromodulation on the underlying perceptual process. The two responses in our task provided a within-trial comparison, since both required a prompt-guided button press under the same sonication condition but engaged different cognitive processes.

In our study, thalamic LIFU did not uniformly speed or slow responses. Its effect was observed for subjective report latency (RT2), but not for object categorization (RT1), and depended on both the acoustic parameters and the focal position of the acoustic field.

In a sequential task, perceptual evidence can continue to accumulate after an initial choice and inform a later judgment [21,22]. By the time the RT2 prompt appeared, subjects had viewed the stimulus, completed the 2-s delay, and made the categorization choice. The selective RT2 interaction therefore implies that LIFU can alter post-categorization processing in the evaluation of perceptual evidence and its conversion into a subjective report.

Among the four intended Tian Scale-II targets, VP was the only target that showed significant DC70–DC5 contrast. We previously found that this region showed predominantly unimodal functional connectivity, with strong somatomotor coupling [17]. This connectivity suggests that motor-related thalamocortical circuitry may contribute to translating a subjective judgment into a manual report.

Beyond the four intended targets, our spatial analysis identified two thalamic voxel sets in which LIFU effect differed between DC conditions (Table 1). The two sets might reflect different components in reporting of conscious experience: perceptual evaluation in posterior pulvinar circuitry and the response selection and execution in anterior mediodorsal–motor circuitry.

The largest components of the anterior voxel set were the parvocellular mediodorsal, ventrolateral posterior, and ventral anterior nuclei. The mediodorsal thalamus integrates task information at the time of decision and response selection through functional coupling with the prefrontal cortex [11,23,24]. On the other hand, it also includes various components of motor thalamus [25]. Ventral anterior and ventrolateral nuclei connect basal ganglia and cerebellar outputs with premotor and supplementary motor cortices [26–28]. The anterior set may therefore encompass multiple circuits involved in selecting a report and translating it into action.

The posterior voxel set was concentrated in the medial and lateral pulvinar. The pulvinar regulates information transmission among visual cortical areas according to attentional demands [29–31]. In a visual confidence task, pulvinar inactivation changed opt-out behavior without impairing categorization performance [32], relevant to our task structure distinguishing between the initial categorization and the subsequent subjective report. Thus, the pulvinar-dominant posterior set may include pathways that coordinate visual information and evaluate the perceptual evidence that informs the subjective report.

The opposing DC dependence of the two territories (Table 1) suggests a possible physiological account. Previous studies have proposed predominantly facilitatory effects of high-DC sonication and suppressive effects of low-DC sonication in human thalamus [33–38]. Under this account, facilitation of anterior mediodorsal-motor circuitry under DC70 would shorten report latency, whereas facilitation of posterior pulvinar-dominant circuitry would prolong visual-attentional evaluation, and suppression under DC5 would reverse these relationships. Perturbation studies provide directionally consistent evidence. Electrical stimulation of mediodorsal and motor thalamus shortened visuomotor reaction times [39,40], while chemically inactivating them lengthened visual task latency [41,42]. Pulvinar perturbation showed the opposite trend. Stimulating it delayed saccadic responses [43], whereas inactivation shortened visual reaction time [44,45].

A methodological contribution of this study is the use of simulated acoustic fields in addition to predefined target labels, extending our previous field-based approach linking perceptual sensitivity to the core–matrix cell gradient [17]. Target-level analyses assign all sonications directed at a target to the same anatomical category. Such analyses do not retain differences in the beam direction, focal position, and intensity distribution across sonications, which are typically treated as targeting error. However, these variations can be informative in thalamic neuromodulation because small anatomical and functional units are embedded within the cytoarchitectural, functional, and connectivity gradients [8,9,46,47]. In our spatial analysis pipeline, we utilized focal spot coordinates and voxel-wise intensity and demonstrated that acoustic field simulations can serve as anatomical variables rather than only as measures of targeting accuracy.

Our study has limitations. First, both DC70 and DC5 were active conditions. The study did not include a sham condition with identical procedures and transducer placement but no ultrasound emission. We therefore cannot determine whether either acoustic protocol lengthened or shortened absolute RT2. Second, by matching temporal-average intensity, the two configurations differed in pulse-average intensity, pressure, and mechanical index. Our interpretations on DC-dependent bidirectional neural effects therefore remain provisional because the two configurations differed not only in DC. Larger pulse-average intensity at DC5 created larger transducer noise and nonspecific auditory confounds could have acted differentially [48]. Third, spatial analyses used acoustic fields simulated on pseudo-anatomical models rather than those measured in vivo. Finally, no functional imaging was conducted, and we cannot directly attribute the observed sonication effects to neural excitation or suppression.

In conclusion, thalamic sonication was associated with the latency of the reporting of conscious visual experience, but not the preceding categorization response. Its effect was both target- and acoustic-protocol-dependent. Under DC70, stronger intensity shortened report latency in anterior thalamus and prolonged latency in posterior thalamus. Both associations were reversed under DC5. Analyzing the sequential reaction times separately, and treating acoustic field variations as an anatomical variable, our work revealed behavioral effects of human LIFU neuromodulation that a single reaction time endpoint and predefined target labels would not fully capture.

## METHODS

### Ultrasound protocol

LIFU was delivered using a BrainSonix BXPulsar 1002 system with a single transducer having 650-kHz fundamental frequency and 80.7-mm free-field focal depth. The transducer was coupled to the right temple and fixed with fabric bands. For neuronavigation, we generated pseudo-anatomical images by linearly rescaling the ICBM 2009c Nonlinear Asymmetric template to match the width, depth, and top-to-ear distance of the subject. BrainSight system was used for neuronavigation.

Two acoustic configurations used in the study (DC70 and DC5) had the same 10-Hz PRF and derated spatial-peak temporal-average intensity (*I*_SPTA.3_) of 0.72 W/cm^2^ (non-derated *I*_SPTA.0_ = 1.02 W/cm^2^), and differed in DC and pulse-average intensity. DC70 used 70-ms pulse width (70% DC) yielding derated *I*_SPPA.3_ = 1.03 W/cm^2^. DC5 used 5-ms pulse width (5% DC) yielding *I*_SPPA.3_ = 14.4 W/cm^2^. Mechanical indices were 0.20 and 0.77, respectively.

Pseudo-anatomical images were segmented using Complete Head Anatomy Reconstruction Method [49] with fat suppression algorithm [50]. Acoustic fields were simulated using the recorded transducer position and angle and normalized to MNI space. Simulated fields were in a 2-mm isotropic grid.

### Thalamic atlas and analysis mask

For LIFU stimulation, four target regions in the left thalamus were selected from Tian Subcortical Atlas Scale-II [8]: ventroanterior (VA, centroid MNI –8, –11, 6 mm), ventroposterior (VP, –13, –23, 2 mm), dorsoanterior (DA, –12, –23, 12 mm), and dorsoposterior (DP, –16, –31, 2 mm). These parcel labels are derived from functional connectivity gradient mapping and are not equated with similarly named anatomical structures.

The thalamic search mask was created as the union of Tian and Morel thalamic atlases [8,18]. From Tian Scale-II atlas, eight thalamic parcels were selected: bilateral VA, VP, DA, and DP. The original Morel atlas comprised 76 hemisphere-specific regions (38 bilateral pairs). Before forming the union, we excluded the bilateral red nucleus, subthalamic nucleus, mammillothalamic tract, and habenula, leaving 68 thalamic nuclei or nuclear subdivisions (34 pairs). Both atlases were represented on the same 1-mm MNI grid.

### Subjects

The parent study was approved by the Institutional Review Board of the University of Michigan Medical School [17]. All subjects provided written informed consent and were compensated. Sixty healthy adults were enrolled (mean age = 25.9 ± 6.3 years). The sample included 38 women and 22 men. All were right-handed, with no hearing loss or color blindness, and had normal or corrected-to-normal vision. Subjects were randomized to DC5 or DC70 (*n* = 30 per arm). Six were excluded due to various technical issues. One completed a different task, three had technical beam targeting problems, one could not complete threshold determination, and one had a Base-1 hit rate performance below 15%. The reaction-time analysis therefore included 54 subjects (27 per arm).

### Trial structure and reaction time outcomes

Each block contained 100 trials displaying 80 real and 20 phase-scrambled images. Each trial began with a fixation interval of 3–6 s sampled from an exponential distribution. The image was presented for 67 ms, followed by a 2-s post-stimulus delay. Subjects then answered two questions, each available for up to 3 s. The first question (Q1) was a four-alternative forced-choice categorization of the stimulus as a face, house, object, or animal. Subjects were instructed to guess even without recognition. The second prompt (Q2) asked whether they had “a meaningful visual experience of the object” (YES/NO). RT1 and RT2 were measured from the display of Q1 and Q2 prompts, respectively, to the subject’s response. If the subject did not respond within the 3-s window, we skipped to the next step, and the corresponding reaction time was saved as N/A.

### Experimental design

Before the main task, a 60-trial adaptive staircase selected image contrast to produce approximately 50% YES responses to Q2 [17]. As a main task, subjects underwent six 100-trial blocks consisting of an initial no-LIFU baseline (Base-1), four LIFU-ON blocks, and a final no-LIFU baseline (Base-2). During the four LIFU-ON blocks, four thalamic targets were stimulated with the fixed DC. The sonication order across the four regions was pseudo-randomized and counterbalanced. Each LIFU-ON block contained twelve 30-s sonication epochs alternating with 30-s intervals, producing 23 alternating ON/OFF epochs over approximately 11.5 min. LIFU epochs were not synchronized to task trials.

### Reaction time preprocessing

RT1 and RT2 were analyzed separately. Seven blocks across five subjects were excluded due to technical problems. From each block, we excluded the initial 10 trials to reduce transient effect (Fig. S1). Trials were included only when the subject responded to both questions, ensuring that two reaction time outcomes used the same paired-valid trial set. Each outcome was summarized as the median across eligible trials.

### Primary behavioral models

We fit separate LME models to block-median RT1 and RT2 with subject-specific random intercepts and order slopes:

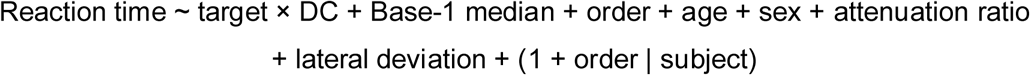

Subject-specific Base-1 median was added as a subject-level covariate to account for individual differences in baseline motor performance. Order denotes the ordinal position (1–4) of the target region within each subject’s four active blocks. Order slopes were added to account for subject-specific task adaptation. Attenuation ratio was the simulated tissue-to-water acoustic-energy ratio at the plane of maximum tissue pressure, expressed in decibels. Lateral deviation was the Euclidean distance in millimeters between the neuronavigation beam vector and the assigned target coordinate [17].

Models were estimated by restricted maximum likelihood. Target-by-DC interactions were evaluated using Satterthwaite *F* tests, with FDR correction across RT1 and RT2 using the Benjamini-Hochberg procedure. For post-hoc testing, two-sided Satterthwaite-adjusted DC70– DC5 contrasts were estimated within each target and corrected across the four contrasts. All statistical tests were two-sided.

### Focal spot analysis

The analysis included 179 simulated LIFU intensity fields and corresponding reaction time outcomes from 47 subjects. Target, DC, and their interaction were excluded:

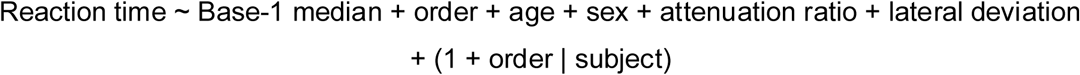

The model was estimated by restricted maximum likelihood. Baseline-adjusted reaction time was defined as the conditional residual after subtracting the fitted fixed effects and subject-specific random intercepts and order slopes.

For each acoustic field, we calculated the 3D MNI coordinates of its focal spot, defined as the 2-mm voxel showing highest simulated intensity. Each coordinate was correlated with baseline-adjusted reaction time using Spearman’s *ρ* separately within DC70 and DC5. Correlations were Fisher-transformed, and the between-DC contrast was defined as Δ*z* = *z*_DC70_ – z_DC5_. We tested each axis using 10,000 subject-level arm-label permutations that kept subject label of each field and preserved arm sizes. Two-sided permutation *p*-values were FDR-corrected across the three axes.

### Voxel-level intensity correlation analysis

The same 179 acoustic fields and baseline-adjusted reaction time values obtained from focal spot analysis were used. A voxel of the simulated intensity field was included only when its entire 8-mm^3^ volume fell within the atlas union (see Methods: Thalamic atlas and analysis mask), yielding 1,923 voxels for analysis. At each voxel, raw simulated intensity was correlated with adjusted reaction time values using Spearman’s *ρ*, separately within DC70 and DC5. All fields were included, including those with zero intensity at that voxel. Correlations were Fisher-transformed to obtain *z*_DC70_ and *z*_DC5_. The between-DC contrast was defined as Δ*z* = *z*_DC70_ – z_DC5_. To assess spatial gradients, we linearly regressed voxel-wise *z*_DC70_, *z*_DC5_, and Δ*z* values against MNI *x*, *y*, and *z* coordinates separately.

For *z*_DC70_ and *z*_DC5_, adjusted reaction time values were randomly reassigned among each participant’s available target-specific acoustic fields. Acoustic fields, target labels, and participant membership remained fixed. We recalculated the voxel-wise Spearman correlations, Fisher-*z* maps, and three spatial slopes for each of 10,000 permutations. For Δ*z*, DC labels were permuted at the participant level while keeping all fields from each participant together. The two condition-specific maps, Δ*z* map, and three spatial slopes were recalculated for each of 10,000 permutations. For slope analysis, permutation *p*-values were FDR-corrected across the nine slope tests. For voxel-wise intensity association, FDR correction was performed across the 1,923 voxels in each map.

### Anatomical composition analysis

Voxels that survived FDR correction for Δ*z* were divided into negative-Δ*z* and positive-Δ*z* sets. For each set, their anatomical distributions were characterized using the Tian Scale-III and Morel atlases. The Tian Scale-III atlas included bilateral superior ventroanterior thalamus, inferior ventroanterior thalamus, medial ventroposterior thalamus, lateral ventroposterior thalamus, medial dorsoanterior thalamus, lateral dorsoanterior thalamus, and dorsoposterior thalamus. We calculated the physical overlap between each native 2-mm isotropic analysis voxel and the atlas regions defined on 1-mm isotropic grids. When an analysis voxel overlapped more than one atlas region, its volume was apportioned according to the overlap with each region. Anatomical composition was expressed as the percentage of the total voxel set volume. Regions contributing less than 1% were combined as Other. Volumes outside the respective atlas were retained as separate non-Tian or non-Morel categories. Morel nuclei were grouped according to the anterior, medial, lateral, and posterior organization of the atlas [19,51].

### Software

The near-threshold task used a custom-modified version of the publicly-available code (https://github.com/BiyuHeLab/NatCommun_Levinson2021/) using the Psychophysics Toolbox v3.0.19.4 (http://psychtoolbox.org/) on MATLAB R2023b. Neuronavigation used proprietary BrainSight software (https://www.rogue-research.com/). Head segmentation used the SimNIBS v4.1.0 (https://simnibs.github.io/simnibs/). Acoustic simulation was performed with BabelBrain v0.4.3 (https://proteusmrighifu.github.io/BabelBrain/) [52]. Behavioral and spatial analyses were performed in MATLAB R2024b and SPM12. Morel atlas characterization was performed in Python.

## ACKNOWLEDGEMENTS

We thank Amy McKinney and Aaron Ellis for their invaluable effort as study coordinators. This work was supported by the National Institute of General Medical Sciences of the National Institutes of Health (grant R01GM103894; PIs: A.G.H. and Z.H.) and the Center for Consciousness Science, Department of Anesthesiology, University of Michigan Medical School, Ann Arbor, Michigan, USA. The content is solely the responsibility of the authors and does not necessarily represent the official views of the National Institutes of Health.

## AUTHOR CONTRIBUTIONS

**Hyunwoo Jang:** Conceptualization, Methodology, Software, Validation, Formal analysis, Investigation, Data curation, Writing – Original Draft, Writing – Review & Editing, Visualization; **Jiyang Liu:** Formal analysis; **Anthony G. Hudetz:** Writing – Review & Editing, Funding Acquisition; **Zirui Huang:** Methodology, Validation, Investigation, Writing – Review & Editing, Supervision, Funding acquisition.

