## Supplementary for "Low-intensity focused ultrasound pulsation along the anterior–posterior thalamic axis differentially modulates the latency of reporting conscious visual experience"

#### SI Figures

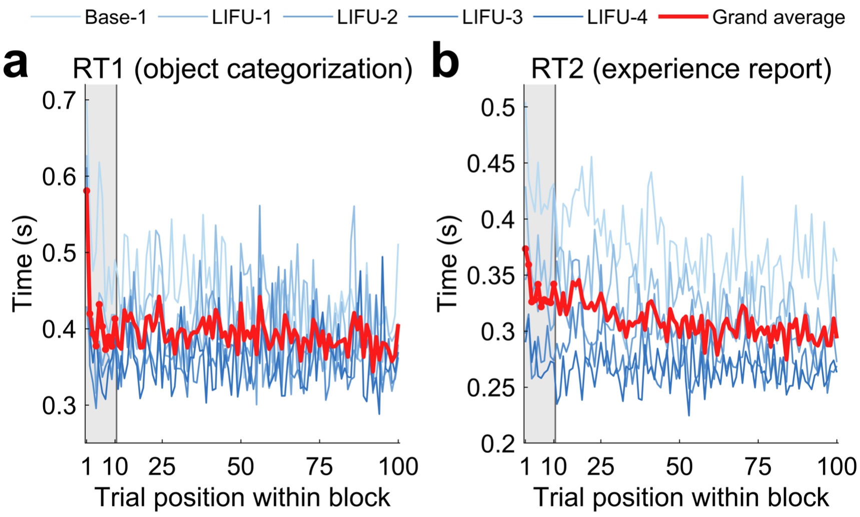


**Fig. S1: Within-block reaction-time trajectories.** Trial-position medians for (a) RT1 and (b) RT2 across subjects. Blue lines show Base-1 and four successive LIFU-ON blocks. Red lines are the mean curves of the five blocks. Gray shading marks trials 1–10 excluded from analysis. RT, reaction time.

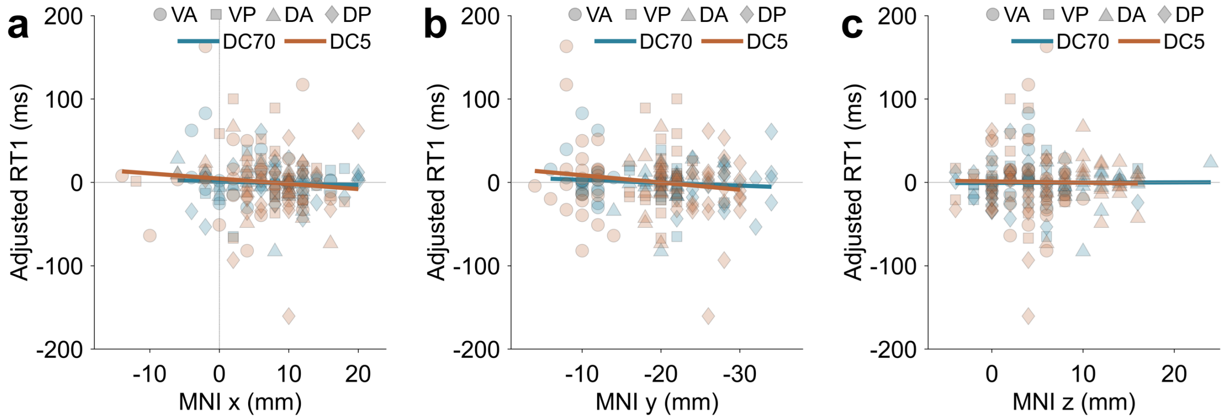


**Fig. S2: Focal spot position and baseline-adjusted RT1.** Baseline-adjusted RT1 are plotted against the MNI (a) *x*, (b) *y*, and (c) *z* coordinates of each simulation field's focal spot. Colors indicate DC and marker shapes indicate target. None of the three between-DC differences survived FDR correction (all $p_{FDR} > 0.05$). DA, dorsoanterior thalamus; DC, duty cycle; DP, dorsoposterior thalamus; RT, reaction time; VA, ventroanterior thalamus; VP, ventroposterior thalamus.

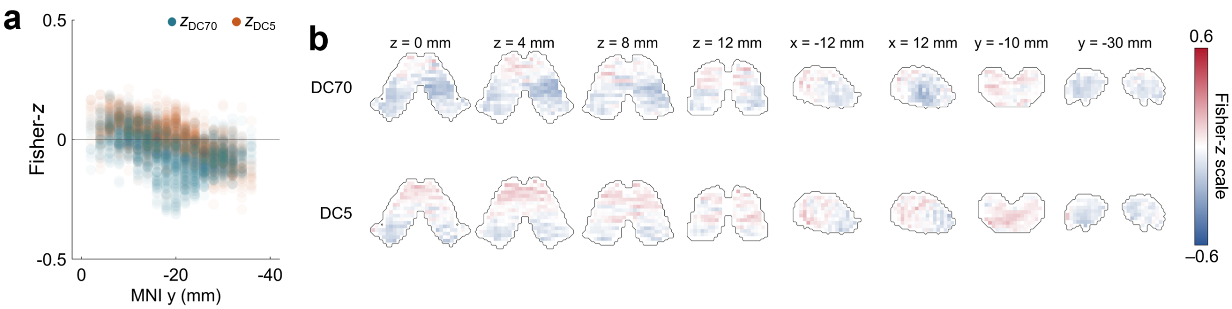


**Fig. S3: Voxel-wise associations between acoustic intensity and categorization reaction time (RT1).** (a) Fisher-transformed Spearman correlations between simulated intensity and baseline-adjusted RT1 plotted against MNI *y* coordinates for each voxel. (b) Same data shown in axial, sagittal, and coronal sections. Outlines indicate the boundary of the Tian S2–Morel union thalamic mask. Corresponding RT2 analyses are shown in Fig. 4. DC, duty cycle; RT, reaction time.

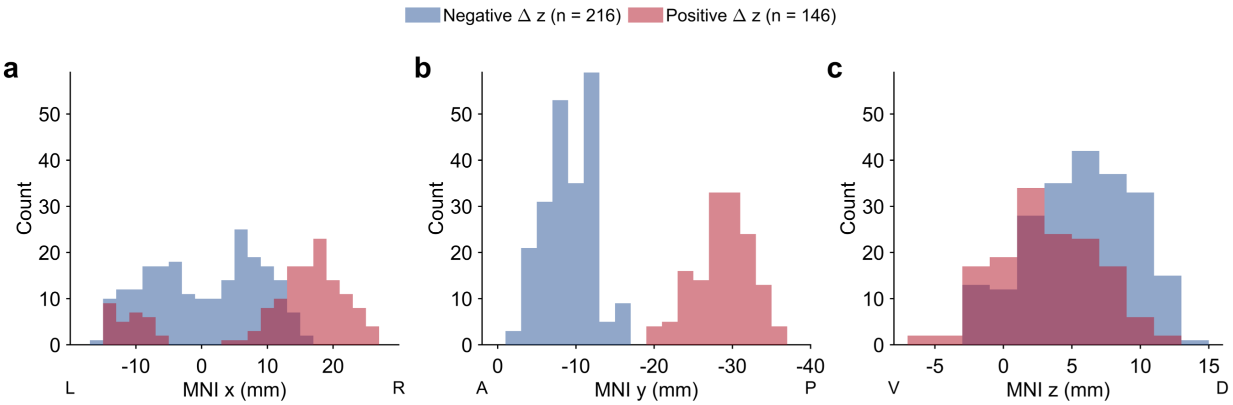


**Fig. S4: Spatial distributions of voxels showing significant between-DC contrasts.** Histograms showing the (a) MNI *x*, (b) *y*, and (c) *z* coordinates of FDR-surviving voxels with negative Δ*z* ($n$ = 216) or positive Δ*z* ($n$ = 146) from Fig. 4e. DC, duty cycle

### SI Tables

##### Table S1: Target-by-DC omnibus tests for reaction time outcomes.

| Outcome | $F$(df1, df2) | Uncorrected $p$ | $p_{FDR}$ |
|---|---|---|---|
| RT1 | 0.730 (3, 121.65) | 0.5361 | 0.5361 |
| RT2 | 4.258 (3, 133.69) | 0.0066 | 0.0132 |

Satterthwaite degrees of freedom were used. $p_{FDR}$ values were corrected across RT1 and RT2.

**Table S2: Target-specific DC70-minus-DC5 contrasts for RT2.**

| Target | Difference (ms) | 95% CI (ms) | Uncorrected $p$ | $p_{FDR}$ |
|---|---|---|---|---|
| VA | 5.4 | [−35.9, 46.7] | 0.7946 | 0.7946 |
| VP | 55.9 | [14.5, 97.3] | 0.0088 | 0.0353 |
| DA | 41.3 | [−0.2, 82.8] | 0.0513 | 0.0684 |
| DP | 45.5 | [4.3, 86.7] | 0.0308 | 0.0616 |

Positive differences indicate longer RT2 under DC70. $p_{FDR}$ values were corrected across the four targets.

**Table S3: Axis-specific associations between focal spot and baseline-adjusted RT2.**

| Axis | $\rho_{DC70}$ | $\rho_{DC5}$ | Δ Fisher-*z* | Uncorrected *p* | $p_{FDR}$ |
|---|---|---|---|---|---|
| Left–right (*x*) | 0.0354 | –0.0747 | 0.1102 | 0.2318 | 0.3477 |
| Anterior-posterior (*y*) | –0.1830 | 0.2847 | –0.4779 | 0.0047 | 0.0141 |
| Dorsal–ventral (*z*) | –0.0272 | 0.1013 | –0.1289 | 0.3813 | 0.3813 |

Within-DC associations are Spearman correlations. Δ Fisher-*z* = atanh($\rho_{DC70}$) − atanh($\rho_{DC5}$). Uncorrected *p* is based on 10,000 subject-level arm-label permutations. $p_{FDR}$ values were corrected across the three axes.

**Table S4: Whole-mask spatial gradients in voxel-wise acoustic intensity–RT2 associations.**

| Condition | *n* | Axis | Slope (Fisher-z/mm) | 95% CI | Uncorrected *p* | $p_{FDR}$ |
|---|---|---|---|---|---|---|
| DC70 | 22 | Left–right (*x*) | 0.0005 | [–0.0034, 0.0040] | 0.8535 | 0.8535 |
| | | Anterior-posterior (*y*) | –0.0141 | [–0.0297, 0.0029] | 0.1106 | 0.2488 |
| | | Dorsal–ventral (*z*) | –0.0013 | [–0.0167, 0.0129] | 0.7981 | 0.8535 |
| DC5 | 25 | Left–right (*x*) | –0.0032 | [–0.0070, 0.0003] | 0.0716 | 0.2148 |
| | | Anterior-posterior (*y*) | 0.0215 | [0.0057, 0.0370] | 0.0090 | 0.0405 |
| | | Dorsal–ventral (*z*) | 0.0074 | [–0.0062, 0.0216] | 0.2686 | 0.4029 |
| DC70–DC5 | 47 | Left–right (*x*) | 0.0037 | [–0.0015, 0.0089] | 0.1808 | 0.3254 |
| | | Anterior-posterior (*y*) | –0.0358 | [–0.0591, –0.0124] | 0.0024 | 0.0216 |
| | | Dorsal–ventral (*z*) | –0.0087 | [–0.0279, 0.0101] | 0.3808 | 0.4896 |

Slopes were estimated separately along each axis across 1,923 thalamic voxels. Confidence intervals were derived from 10,000 permutation distributions. $p_{FDR}$ values were corrected across the nine slope tests. Positive slopes indicate increasing Fisher-*z* values toward the right, anterior, or dorsal direction.
